# Data mining antibody sequences for database searching in bottom-up proteomics

**DOI:** 10.1101/2024.02.13.580076

**Authors:** Xuan-Tung Trinh, Rebecca Freitag, Konrad Krawczyk, Veit Schwämmle

## Abstract

Mass spectrometry (MS)-based proteomics allows identifying and quantifying thousands of proteins but suffers from challenges when measuring human antibodies due to their vast variety. The mainly used bottom-up proteomics approaches rely on database searches that compare experimental values of peptides and their fragments to theoretical values derived from protein sequences in a database. While the human body can produce millions of distinct antibodies, the current databases for human antibodies such as UniProtKB/Swiss-Prot are limited to only 1095 sequences (as of 2024 Jan). This limitation may hinder the identification of new antibodies using mass spectrometry. Therefore, extending the database for mass spectrometry is an important task for discovering new antibodies. Recent genomic studies have compiled millions of human antibody sequences publicly accessible through the Observed Antibody Space (OAS) database. However, this data has yet to be exploited to confirm the presence of these antibodies. In this study, we adopted this extensive collection of antibody sequences for conducting efficient database searches in publicly available proteomics data with a focus on the SARS-CoV-2 disease. Thirty million heavy antibody sequences from 146 SARS-CoV-2 patients in the OAS database were digested *in silico* to obtain 18 million unique peptides. These peptides were then used to create new databases for bottom-up proteomics. We used those databases for searching new antibody peptides in publicly available SARS-CoV-2 human plasma samples in the Proteomics Identification Database (PRIDE). This approach avoids false positives in antibody peptide identification as confirmed by searching against negative controls (brain samples) and employing different database sizes. We show that the found sequences provide valuable information to distinguish diseased from healthy and expect that the newly discovered antibody peptides can be further employed to develop therapeutic antibodies. The method will be broadly applicable to find characteristic antibodies for other diseases.

## Introduction

The human antibody, an essential immune system component, is a protein molecule composed of two heavy and two light polypeptide chains interconnected by disulfide bonds [1]. They are Y-shaped proteins produced by B cells in response to the presence of foreign substances called antigens [2]. Antibodies could be used as biomarkers for diagnostics of various infectious, autoimmune, and oncological diseases [3,4]. Moreover their chemical flexibility makes them an asset for identification, isolation, and labeling of proteins [5–7]. Furthermore, antibodies are increasingly utilized in therapy for treatments of diseases [8]. Functions of antibodies are largely determined by their amino acid sequences.

Mass spectrometry (MS)-based proteomics is an important method for identifying and quantifying antibodies by analyzing the mass of ionized biomolecules [9,10]. Bottom-up tandem liquid-chromatography mass spectrometry (LC-MS) is indispensable to analyze antibodies in intricate mixtures such as blood plasma. In the bottom-up proteomics approach, proteins undergo enzymatic digestion into peptides, and these resulting peptides are subjected to MS and MS/MS analysis for protein identification and quantification [11,12]. For the identification of thousands of peptides, database search is the main method that entails comparing experimental mass of peptides and their fragments obtained from MS and MS/MS with theoretical values derived from protein sequences in a database [11,12]. UniProt is the mainly used protein database, containing 20,422 entries of reviewed human proteins and 1095 entries of antibodies (as of 2024 Jan) [13,14]. These numbers are significantly lower than the expected millions of different antibodies produced by the human body. Such a severely limited number of antibody sequences in current human protein databases poses a challenge for bottom-up proteomics in detecting reasonable numbers of antibodies within complex mixtures, such as blood plasma.

Recent genomic studies have compiled millions of potential human antibody sequences publicly accessible through the Observed Antibody Space (OAS) database [15,16]. Such a resource poses a potential database for antibody detection. We show that including this OAS data to the database search allows the detection of previously undetected antibody peptides. We explore how the extensive collection of antibody sequences in the OAS database enriches database searches of proteomics data. The newly discovered antibody peptides in the SARS-COV-2 samples are further assessed for the diagnostic and therapeutic value.

## Methods

The workflow of this study contains four main steps: antibody data collection, peptide data pre-processing, database creation, and proteomics database search (Fig. 1).

**Fig. 1.**
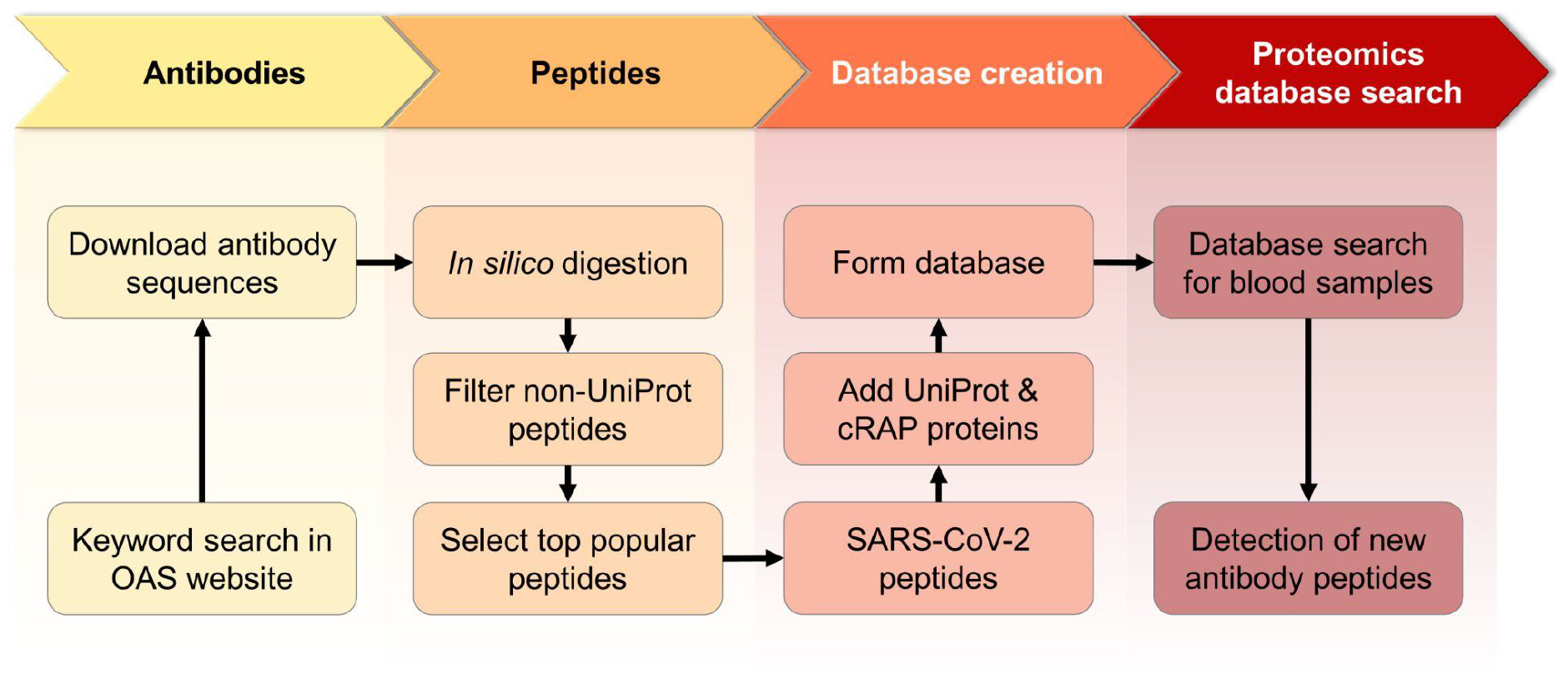
Workflow of the study

### Antibody data collection

The Observed Antibody Space database (OAS) [15,16] constitutes a comprehensive endeavor aimed at assembling and annotating immune repertoires tailored for extensive-scale analysis. It currently contains over two billion sequences, from 90 different studies. These repertoires include a wide array of immune states, primarily focusing on human and mouse organisms and individual variations. The dataset has undergone rigorous processes including sorting, cleaning, annotation, translation, and numerical indexing, rendering it meticulously prepared for dissemination online (https://opig.stats.ox.ac.uk/webapps/oas/) for download.

We downloaded SARS-CoV-2 antibody data from the Observed Antibody Space database by using the following keyword search: “Heavy chain”, “SARS-CoV-2”, “BSource of PBMC’’, “undefined longitudinal”, “human species”, “none vaccine”, and “defined subject”. The search was conducted on 10th February 2023, resulting in 30,966,193 heavy-chain antibody sequences. This dataset was denoted as OAS. Next, we downloaded human protein data from the UniProt database by using the keyword search “human” and choosing the “Reviewed (Swiss-Prot)” and “canonical & isoform” options. The search was conducted on 07th March 2023 and gave 42,421 human protein sequences. This dataset was denoted as UniProt.

### Data pre-processing

The UniProt protein and OAS antibody sequences were then digested *in silico* into peptide sequences using Protein Digestion Simulator software [17,18]. The digestion settings were: fully tryptic (KR not P), max missed cleavages of 0, minimum fragment mass of 400, and maximum fragment mass of 6000. The digestion process produced 691,027 and 18,465,822 different peptides of UniProt and OAS, respectively.

For the OAS peptide sequences, we removed the peptides that showed up in UniProt. After this removal, we obtained 18,419,969 distinct peptide sequences which were found in nine different bio-projects [19–27]. Employing a database containing 18 million sequences drastically increases the search space, leading to prolonged search times and considerable difficulties in controlling the false discovery rate. Therefore, among 18 million peptides, we selected those that are commonly present in the highest number of antibodies to create databases of different sizes, aiming to minimize search time and the false discovery rate.

### Databases preparation

The top 10^2^, 10^3^, 10^4^, 10^5^, 10^6^ and 10^7^ peptides which were commonly present in the highest number of antibodies in the OAS data were then combined with UniProt and cRAP protein sequences (contaminant proteins, https://www.thegpm.org/crap/) to form six databases labeled as DB_1-6_ for later database search. The databases are available at Zenodo [28]. Source code for database preparation is available at github (https://github.com/trinhxt/SDU_Immunoinformatics).

### Proteomics database search for SARS-COV-2 antibody peptides

Database search was conducted using FragPipe [29–37] with the above databases and six PRIDE datasets including four blood plasma datasets (PXD031813, PXD029181, PXD023175, and PXD020354) [38–40], two depleted blood plasma datasets (PXD036491 [41] and PXD022296) and one brain dataset (PXD039808) [42]. Depleted blood plasma samples were samples of which most abundant proteins (e.g., antibody, fibrinogen, etc.) were removed. The brain dataset was used as a negative control sample because the presence of blood-brain barrier restricts the movement of antibodies from the bloodstream into the brain tissue, which is why brain samples are generally devoid of antibodies when directly analyzed [43]. Parameter settings for each dataset were adapted from the previous studies that produced the datasets (**Table S1**).

### Utilizing newly found antibody peptides for differentiating patient samples

For distinguishing patient samples (COVID, sepsis, flu, healthy), we developed random forest based classification models using the following two different datasets. In dataset 1, we included known antibody peptides from the UniProt database as features. In dataset 2, we included both known antibody peptides from the UniProt database and newly found antibody peptides from our new databases as features. These features had values of 0 and 1 (i.e., absence and presence of a peptide in samples). In both datasets, the endpoint was category (COVID, healthy, sepsis, and flu), and there were 491 peptide sets corresponding to 491 samples (286 COVID, 135 healthy, 56 sepsis and 14 flu samples). Classification models were developed by using 80% of data for training and 20% for testing. Receiver operating characteristic (ROC) curve and area under the curve (AUC) were used for accessing performance of the models in classification of patient samples.

## Results

First, we investigated how database size influences the detection of antibodies in public data. Using the workflow in the Method section (Fig. 1), we obtained distinct databases containing various numbers of common SARS-CoV-2 antibody peptides (Fig. 2). By testing these databases for proteomic search on data PXD029181, we could find which database was the most suitable for reaching low analysis run times and high numbers of detected peptides (Fig. 3). Next, we utilized the optimized database for proteomic search of other data to see how many new antibody peptides were detected in previously analyzed samples (Fig. 4). Among newly found peptides, we focused on peptides which were in the most variable region of antibody heavy chains (CDR-H3). CDR-H3 peptides were found in all sample groups but some peptides were overrepresented in SARS-CoV-2 samples (Fig. 5-6) and could be used in differentiating SARS-CoV-2 and healthy samples (Fig. 7).

**Fig. 2.**
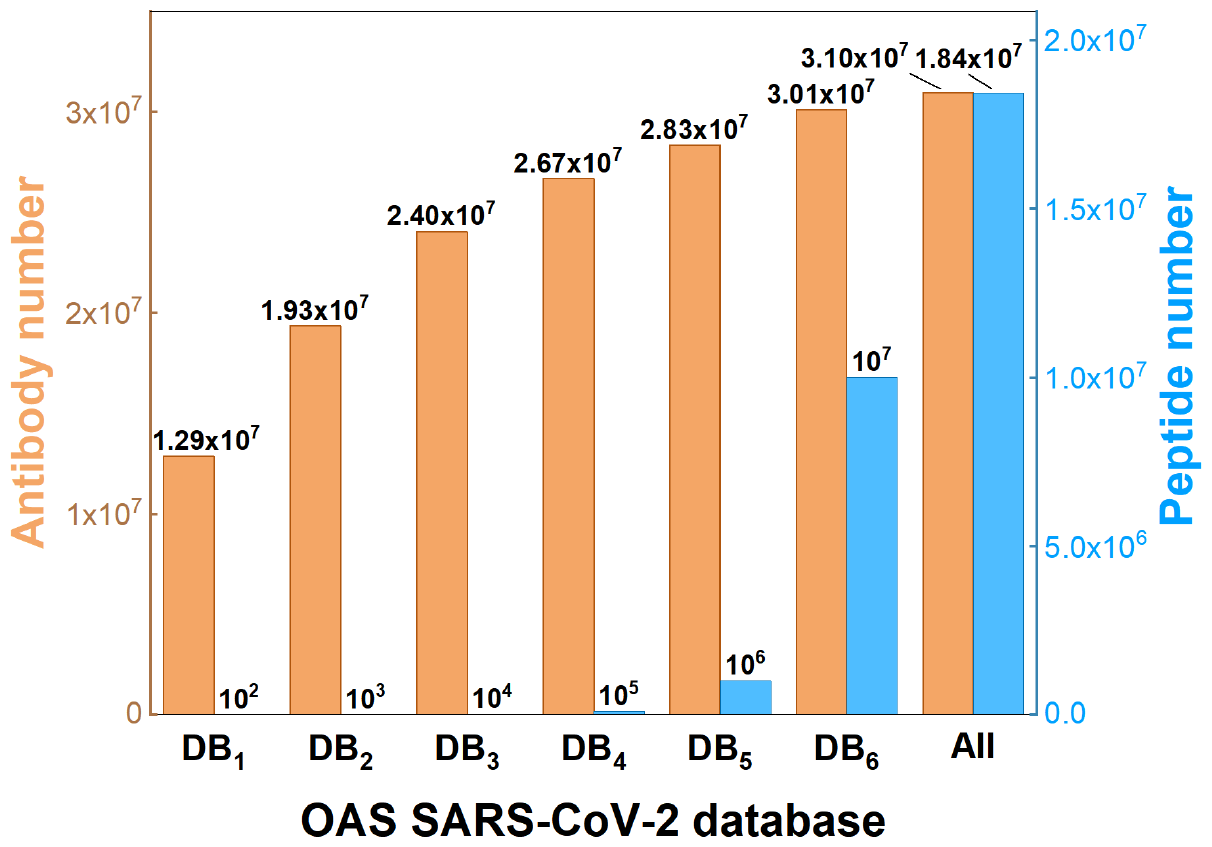
Number of peptides and corresponding antibodies of different databases created by filtering top common peptides in OAS SARS-CoV-2 data.

**Fig. 3.**
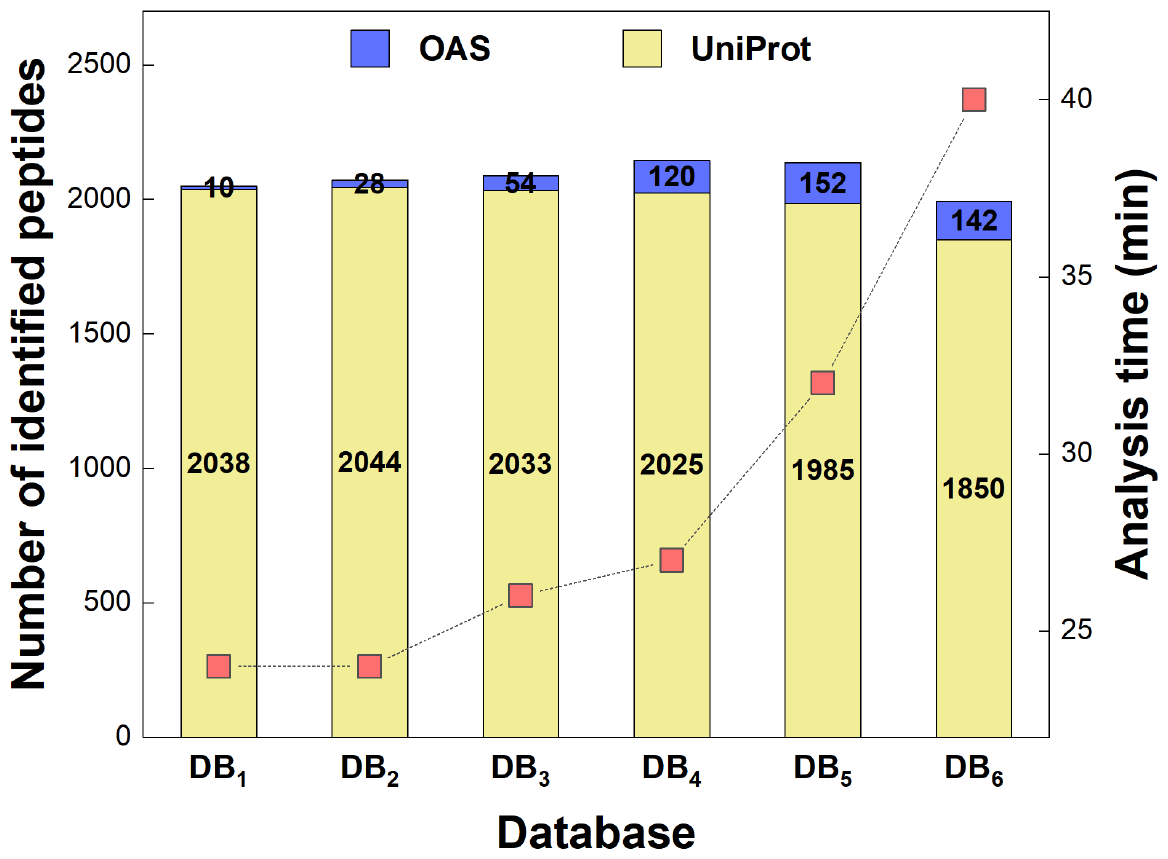
Representative analysis time and number of identified peptides in blood plasma sample Plasma_S86 of PRIDE project PXD029181 when running the database search with different database sizes (DB_1_ - DB_6_). The red squares denote the typical analysis time on a computer with Intel Xeon Gold 6130 processor and 24GB of RAM.

**Fig. 4.**
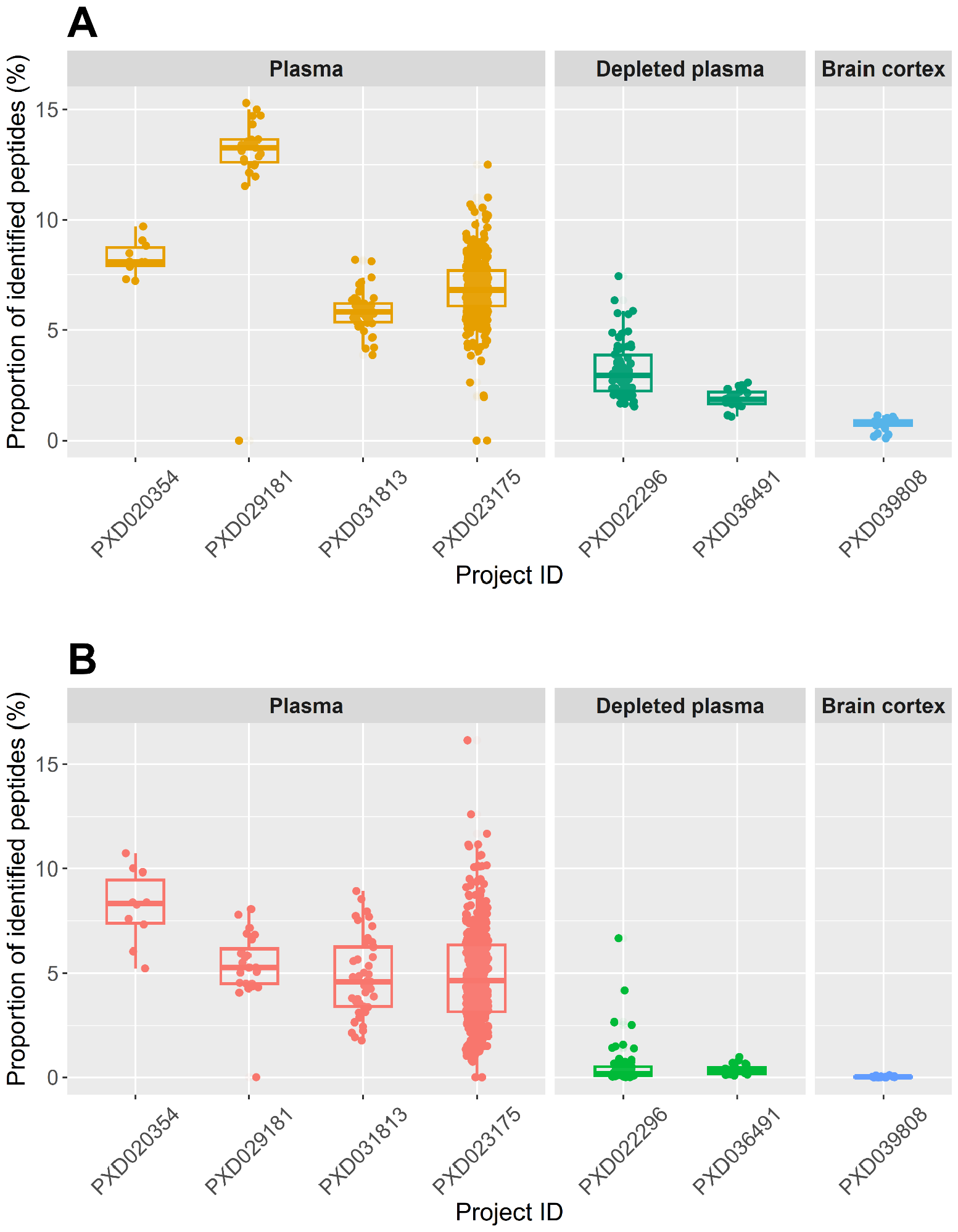
**A)** The proportion of antibody peptides in UniProt relative to the total identified peptides and **B)** the proportion of new antibody peptides relative to the total identified peptides in samples from various PRIDE projects. The database search was conducted using DB_4_.

**Fig. 5.**
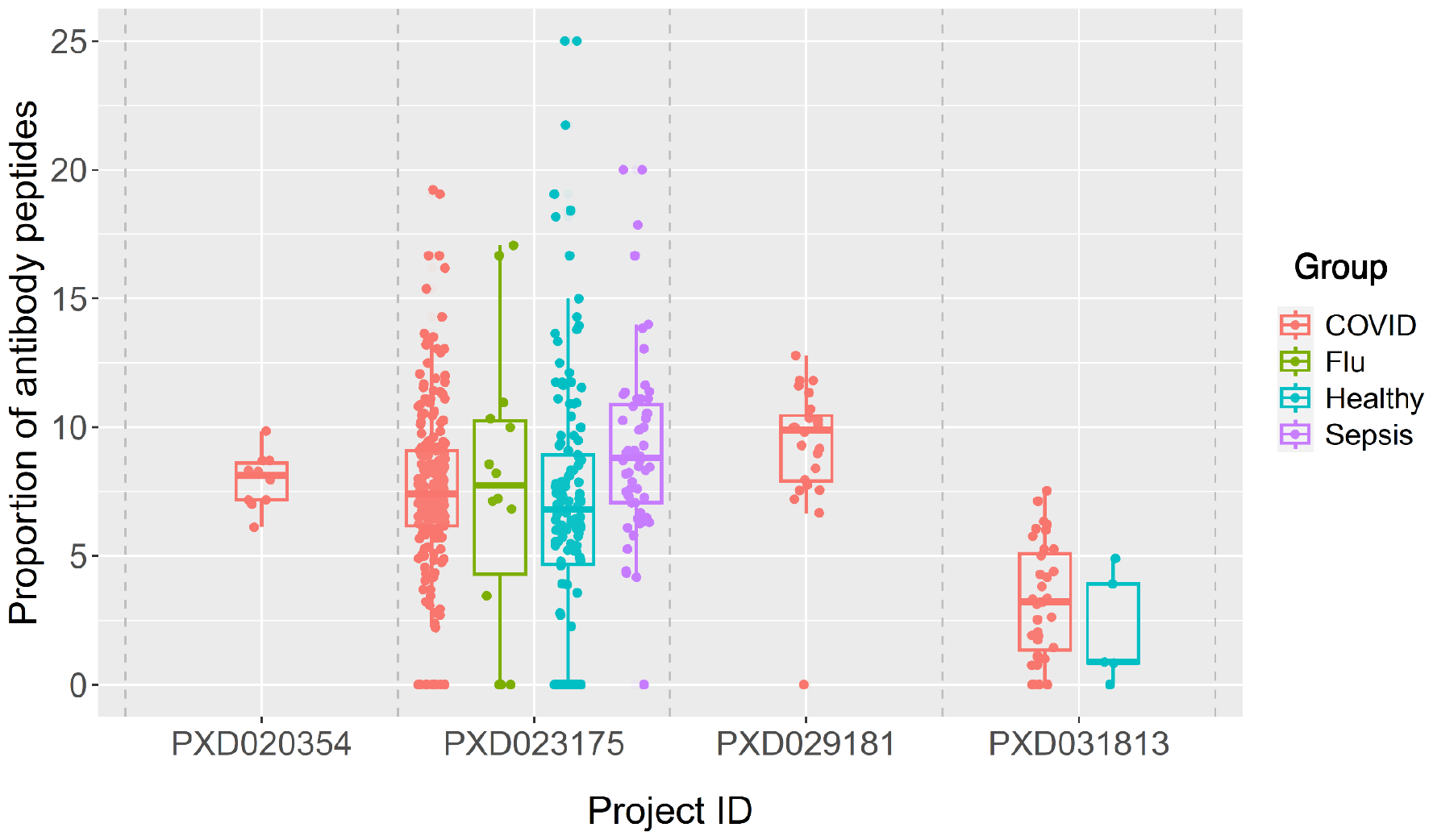
Percentage Of CDR-H3 peptides relative to the total number of antibody peptides identified in all samples.

### OAS-SARS-COV-2 antibody databases show high diversity

After *in silico* digestion of 30,966,193 heavy-chain antibody sequences and removal of identical peptides already present in the human proteome (UniProt human peptides), we remained with 18,419,969 different peptides of OAS data. The number of OAS peptides is smaller than the number of OAS antibodies because many peptides are shared between multiple antibodies.

We compared physicochemical properties of peptides in OAS data to known peptides in the UniProt data for sanity checking (Fig. S1-2). Antibody peptides (OAS) are longer and heavier than UniProt peptides (Fig. S1A-B). UniProt peptides have a higher variety of isoelectric points and charge than the OAS peptides (Fig. S1C-D). The majority of the first amino acids in peptides are glycine, glutamic acid and leucine, while the majority of the second amino acids are leucine, serine, threonine and valine (Fig. S2). The OAS peptides have more C-terminal serine ending than UniProt peptides.

By filtering for the most common peptides in the OAS data, we obtained different datasets having different numbers of OAS peptides and OAS antibody coverages (Fig. 2). DB_1_ - DB_6_ contain top 10^2^ - 10^7^ OAS peptides corresponding to increasing number of antibodies in the OAS data. Increasing the database size to millions of entries inflates the search space leading to very long search times and considerable difficulties with controlling the false discovery rate. Thus, choosing a database that covers large amounts of antibodies and minimizes search time is important. We observed that, for blood sample Plasma_S86 (data PXD029181 in PRIDE database), larger database sizes (DB_1-6_) increase analysis time (up to 24-40 minutes) and the number of detected peptides (Fig. 3). The number of identified UniProt peptides remains relatively consistent across DB_1_ to DB_4_, while there is a significant increase in the number of identified OAS peptides. However, upon expanding the database size to include DB_5-6_, alongside a notable increase in the identification of OAS peptides, we observe a significant decrease in the number of identified UniProt peptides. A Venn diagram shows that identified peptides from larger databases contain the identified peptides from smaller databases (Fig. S3). In what follows, we chose DB_4_ which contains 10^5^ OAS peptides and covers 2.67×10^7^ (86.2%) antibodies of OAS SARS-CoV-2 data to balance analysis time and number of detected peptides.

### Database search consistently finds new antibody peptides

We compared the proportions of known and newly identified antibody peptides in the following three sample groups: blood plasma, depleted blood plasma and brain cortex, to find out whether genuine antibody peptides were detected using our new databases. The amount of detected UniProt and OAS peptides in 491 samples of blood plasma, 100 samples of depleted blood plasma, and 21 samples of brain cortex is shown in Fig. 4. The number of detected UniProt peptides in blood plasma samples (5-15% of detected peptides) is significantly higher than depleted blood plasma samples (2-7% of peptides number) (Fig. 4A). The number of detected UniProt peptides in brain cortex samples is almost zero (average of 0.8% of protein number) (Fig. 4A). Similarly, number of identified OAS peptides in blood plasma samples (1-11% of detected peptides) is also significantly higher than depleted blood plasma samples (0.1-2.5% of detected peptides); and brain cortex samples have almost zero antibody peptides (average of 0.1% of peptide number) (Fig. 4B). This result indicates that by using our created databases, we can detect not only known peptides (UniProt) but also new antibody peptides (OAS SARS-CoV-2). The decreasing number of detected OAS peptides in the depleted blood plasma samples and the near-zero percentage of OAS peptides in the brain cortex samples confirm that the newly detected peptides in the samples are genuine antibody peptides.

### Some CDR-H3 peptides are overrepresented in COVID samples

Antigen binding and specificity of antibodies are significantly influenced by the third complementarity-determining regions of heavy chains (CDR-H3). In this study, identified peptides were mapped back to the CDR-H3 regions in OAS antibody data to determine if they corresponded to CDR-H3 peptides. The proportions of CDR-H3 peptides in blood plasma samples are depicted in Fig. 5. The proportion of CDR-H3 peptides relative to the total number of identified antibody peptides in COVID (SARS-CoV-2) samples is not significantly different from that in healthy samples. Additionally, flu and sepsis samples exhibited slightly higher proportions of CDR-H3 peptides than the healthy samples in Bioproject PXD023175. The overlap among found CDR-H3 peptides in COVID, healthy, flu and sepsis samples is shown in Fig. 6. Among all CDR-H3 peptides, 83 peptides are exclusive to only COVID samples. Healthy and sepsis samples have 6 and 2 specific CDR-H3 peptides, respectively. Flu samples do not have any specific peptides. Lists of these peptides are shown in supporting information. The 83 COVID-specific peptides appeared in 1,274,346 antibodies, while 13 common peptides across all sample groups were found in 529,442 antibodies of the OAS database. Hence, CDR-H3 peptides are considerably overrepresented in COVID samples, also when taking into account the larger number of COVID samples in comparison to healthy and the other diseases. This indicates that the immune system becomes more active after SARS-CoV-2 infection.

**Fig. 6.**
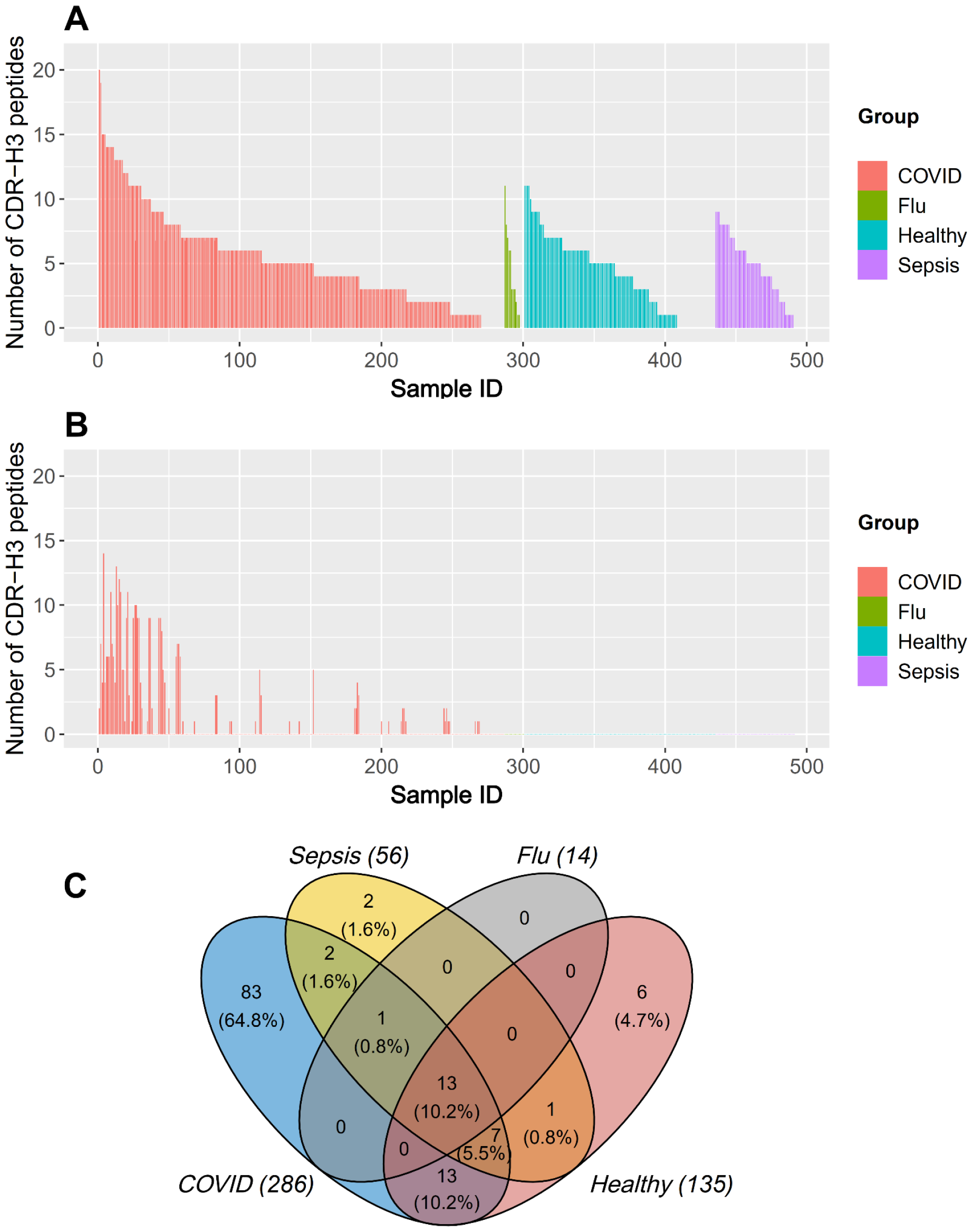
**A)** Distribution of the number of CDR-H3 peptides across samples. **B)** Number of CDR-H3 peptides specific to SARS-CoV-2 in every sample. **C)** Overlap of all identified CDR-H3 peptides across the four groups (COVID - 286 samples, healthy - 135 samples, sepsis - 56 samples, and flu - 14 samples).

**Fig. 7.**
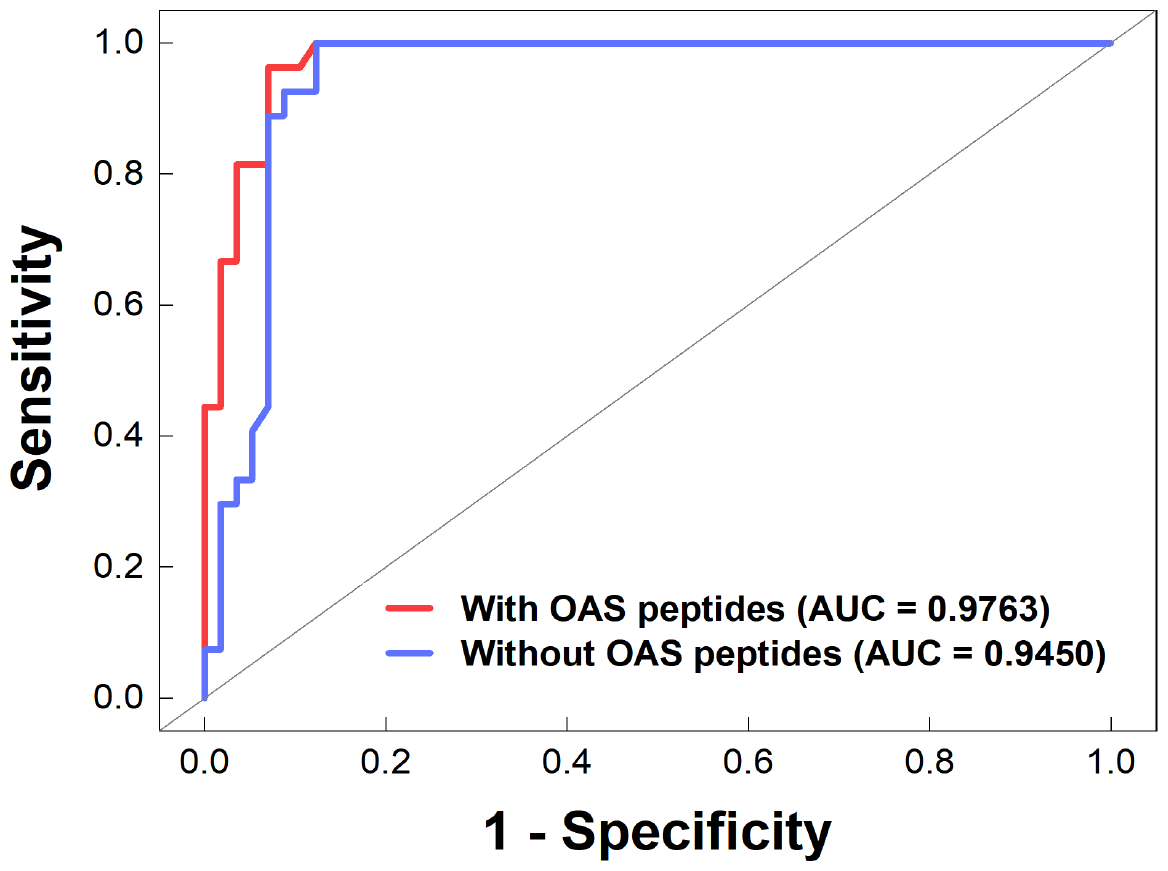
ROC curves of random forest models for classification of COVID and healthy samples using antibody peptides including OAS data (red line) and not including OAS peptides (blue line).

### Newly found antibody peptides improve classification of patient samples

The OAS peptides which were identified in 491 samples were used in classification models of patient samples (COVID, healthy, sepsis, flu). ROC curves and AUC values of two models (one including and one excluding OAS peptides as features) are shown in Fig. 7. The model including OAS peptides shows better classification of COVID and healthy samples (red ROC curve with AUC = 0.9763) than the model not using new OAS peptides (blue ROC with AUC = 0.9450), showing that the found OAS peptides provide relevant information about the disease.

## Discussion

Both next-generation sequencing (NGS) and proteomics play crucial roles in antibody analysis. NGS allows for the deep sequencing of antibody variable regions, providing insights into genetic diversity, clonal expansion, and sequence variations. On the other hand, proteomics focuses on detecting the actual protein products, including antibodies, and enabling the study of expression levels, post-translational modifications, and potential interactions. Proteomics also enables the identification of antibody biomarkers for disease diagnosis [44], monitoring of treatment response [45], and development of personalized medicine approaches [46]. We show that integrating NGS and proteomics data offers insights into active antibody repertoires, bridging genetic information with functional characteristics of antibodies. This integration can be particularly valuable in antibody-based research, such as monoclonal antibody development and understanding immune responses. For example, VanDuijn *et al* combined NGS and proteomics to obtain complementary information in the characterization of the immune repertoire in groups of rats after immunization with purified antigens [47]. Wang *et al* used NGS and proteomics to detect SARS-CoV-2 RNA and proteomic changes in cerebrospinal fluid from COVID-19 patients with neurological manifestations [48].

Inspired by these studies, our work leveraged NGS output for antibody sequencing, constructing novel antibody peptides databases to expand the search space of proteomics database searches while keeping it at a manageable level. Through the utilization of these databases, we did not only re-identify known peptides from UniProt but also discover new antibody peptides. Importantly, these novel peptides were notably absent in negative control samples (brain cortex), showing them to not result from an inflation of false positives.

The discovery of novel antibody peptides offers two promising applications. Firstly, the identified CDR-H3 peptides, specific to COVID samples, hold potential for disease diagnostics. The methodology developed in this study may find broader applicability across various diseases. Secondly, the newly discovered antibody peptides enhance the classification rate between COVID and healthy samples (Fig. 7). This improvement could be leveraged in machine learning models for patient detection using peptides derived from mass spectrometry analysis.

## Conclusion

This study introduces novel antibody databases designed for use in proteomic database searches. The efficacy of these databases was demonstrated through successful testing in both blood plasma and brain cortex samples. Notably, the results indicated the detection of over 5% of previously undiscovered antibody peptides in blood plasma samples. Among these newly identified peptides, approximately 4-10% were found within the CDR-H3 region, with 83 peptides specifically associated with COVID samples. Moreover, the inclusion of new antibody peptides led to an enhanced classification of healthy and COVID samples. Our method is applicable to other diseases and phenotypes, and thus provides a new powerful way for disease characterization, diagnosis and potential treatments.

