## Supplementary for "Data mining antibody sequences for database searching in bottom-up proteomics"

### Supplementary data

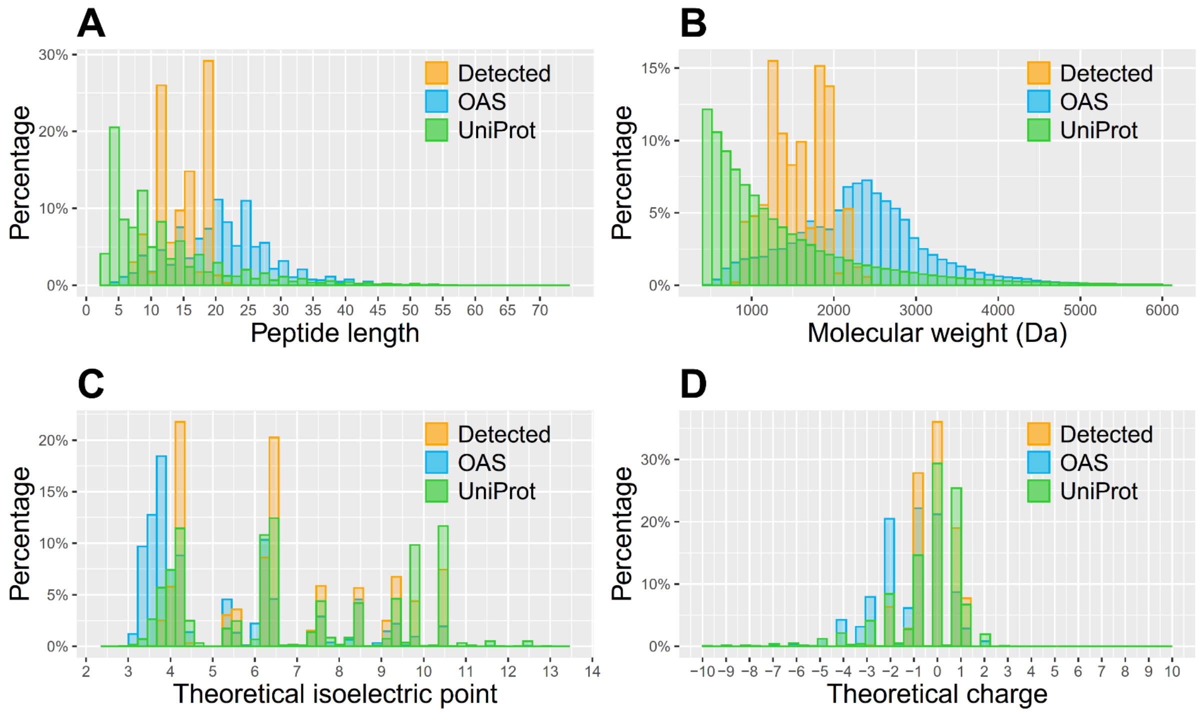


**Fig. S1.** Distribution of (A) peptide length, (B) molecular weight, (C) theoretical isoelectric point, (D) theoretical charge of OAS SARS-COV-2 peptides (N = 18,419,969 peptides), UniProt human peptides (N = 646,863) and new detected peptides in PRIDE projects (N = 3600).

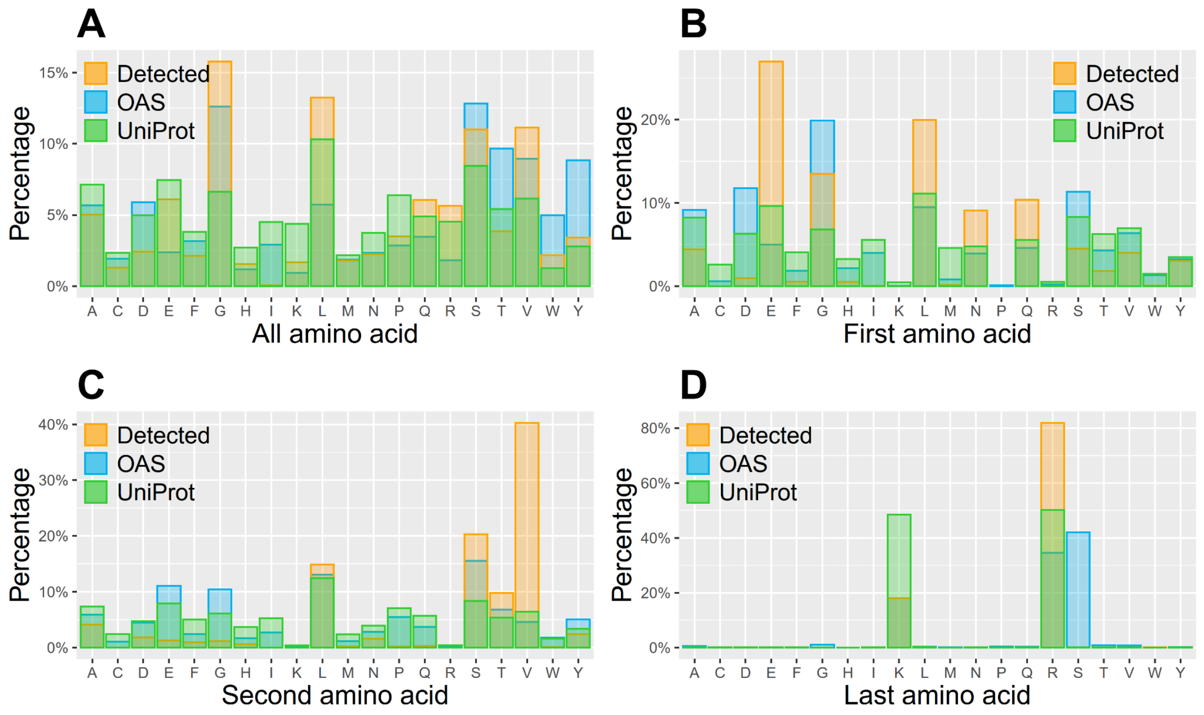


**Fig. S2.** Distribution of the all (A), first (B), second (C), and last (D) amino acids of peptides in OAS-SARS-COV-2 and UniProt data and detected peptides in PRIDE data.

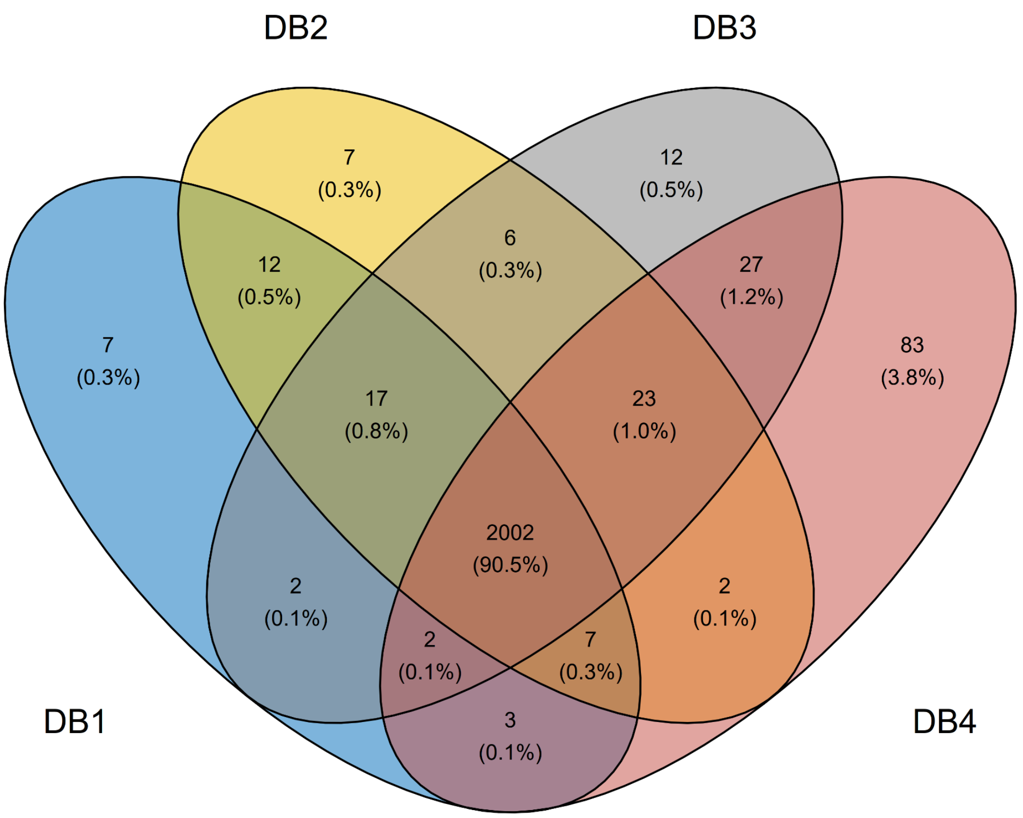


**Fig. S3**. Overlapping identified peptides found in the blood sample Plasma_S86 of PRIDE project PXD029181 by using different databases ($DB_1$ - $DB_4$).

**Table S1**. Parameter settings for database search

| No | PRIDE project | Sample | Missed cleavage | Precursor mass tolerance | Fragment mass tolerance | Fixed modification | Dynamic modifications | FDR |
|---|---|---|---|---|---|---|---|---|
| 1 | PXD031813 | Blood plasma | 2 | ±10 ppm | ±0.02 Da | Cysteine Carbamidomethylation | Phosphorylation and oxidation | .01 |
| 2 | PXD029181 | Blood plasma | 3 | 15.0 ppm | 0.4 Da | Carbamidomethylation: 57.02 | Oxidation (HW) 15.99, Oxidation (M): 15.99, Phosphorylation (STY): 79.97, Deamidation (NQ): 0.98, Acetylation (K): 4.01, Methylation (KR): 14.02 | 0.01 |

| | | | | | | | | |
|---|---|---|---|---|---|---|---|---|
| 3 | PXD020354 | Blood plasma | 2 | 6-ppm | 20 ppm | Cysteine carbamido-methylation | N-acetylation of protein and oxidation of methionine | 0.01 |
| 4 | PXD036491 | Depleted blood plasma | 2 | 10 ppm | 0.02 Da | Cysteine Carbamidomethylation | | 0.01 |
| 5 | PXD022296 | Depleted blood plasma | 2 | 10 ppm | 0.02 Da | Cysteine Carbamidomethylation | Methionine oxidation (+15.994915 Da) and alkylation at N-terminals of protein. | 0.01 |
| 6 | PXD039808 | Brain cortex | 2 | 20 ppm | 0.1 Da | Cysteine carbamidomethylation | Methionine oxidation, asparagine and glutamine deamidation, acetylated N-terminal residues. | 0.01 |
